# Low-input and single-cell methods for Infinium DNA methylation BeadChips

**DOI:** 10.1101/2023.09.18.558252

**Authors:** Sol Moe Lee, Christian E. Loo, Rexxi D. Prasasya, Marisa S. Bartolomei, Rahul M. Kohli, Wanding Zhou

**Affiliations:** Center for Computational and Genomic Medicine, The Children’s Hospital of Philadelphia, PA, 19104, USA; Graduate Group in Biochemistry and Biophysics, University of Pennsylvania, Philadelphia, PA, 19104, USA; Department of Cell and Developmental Biology, Epigenetics Institute, University of Pennsylvania Perelman School of Medicine, Philadelphia, PA 19104, USA; Department of Medicine, University of Pennsylvania, Philadelphia, PA, 19104, USA; Department of Pathology and Laboratory Medicine, University of Pennsylvania, Philadelphia, PA, 19104, USA

## Abstract

The Infinium BeadChip is the most widely used DNA methylome assay technology for population-scale epigenome profiling. However, the standard workflow requires over 200 ng of input DNA, hindering its application to small cell-number samples, such as primordial germ cells. We developed experimental and analysis workflows to extend this technology to suboptimal input DNA conditions, including ultra-low input down to single cells. DNA preamplification significantly enhanced detection rates to over 50% in five-cell samples and ∼25% in single cells. Enzymatic conversion also substantially improved data quality. Computationally, we developed a method to model the background signal’s influence on the DNA methylation level readings. The modified detection *p*-values calculation achieved higher sensitivities for low-input datasets and was validated in over 100,000 public datasets with diverse methylation profiles. We employed the optimized workflow to query the demethylation dynamics in mouse primordial germ cells available at low cell numbers. Our data revealed nuanced chromatin states, sex disparities, and the role of DNA methylation in transposable element regulation during germ cell development. Collectively, we present comprehensive experimental and computational solutions to extend this widely used methylation assay technology to applications with limited DNA.

## INTRODUCTION

DNA modifications, including 5-methylcytosines (5mCs) and 5-hydroxymethylcytosines (5hmCs), are canonical forms of epigenetic modification in human and other mammalian genomes. DNA methylation is found mainly in CpG dinucleotide contexts, where it is extensively implicated in gene transcriptional regulation, cell identity maintenance, organismal development, aging, and diseases (1). Infinium DNA methylation BeadChips are among the most popular genome-wide methylation assays in humans and other species due to the ease of experiment and data analysis (2). These arrays have been the primary data generation workhorse for large data consortia such as The Cancer Genome Atlas (TCGA), with over 160,000 public datasets deposited to Gene Expression Omnibus (GEO) (3). While the adoption of sequencing-based methods is catching up in case and mechanistic studies, the Infinium technology remains the most used assay platform for population-level studies such as meQTL studies (4,5), epigenetic risk scores (6,7), and other epigenome-wide association studies (8,9). In addition to being a powerful and popular tool for biological discovery, the technology has recently enabled rapid clinical application development (10). This platform has found wide success in cancer diagnosis (11), cell-free liquid biopsy (12), and forensics (13). Recently, the Infinium BeadChips have also been used to generate the largest DNA methylome atlas across different mammalian species (14-16).

Despite these successes, a significant drawback of this technology is that it requires over 200 ng of input DNA from the standard processing protocol (2). This requirement constrains scientific and clinical applications with limited DNA availability. For example, as few as 25 primordial germ cells (PGCs) can be found in the early mouse embryo (17). Serum-derived, tumor-originated cell-free DNA (cfDNA) in cancer patients (18) holds value in non-invasive early cancer diagnosis (19). But as little as five ng/ml cfDNA in healthy subjects and 30 ng/ml in cancer patients (20) may be available. These DNA amounts are much lower than the Infinium array standard protocol requires.

In addition to circumstances where the input DNA quantity is limited, it may be of interest to study a complex tissue to dissect cell-to-cell heterogeneity purposefully, even when there is no shortage of total DNA in the tissue. DNA methylation encodes distinctive cell identity fingerprints, which can be used to inform cellular phenotypes (1), trace the state of the DNA-releasing cells (21), and inform cell proportions (22). By performing DNA methylation analysis on laser-capture microdissected specimens, one can compare methylomes at different locations in tumors (23,24) or select specific cell types from the brain for analysis (25). In most cases, laser-capture microdissected tissues are limited in quantity and involve pre-assay whole genome amplification, as is done with SNP arrays (26).

The extreme of increasing the cell resolution at the cost of working with small DNA amounts is epitomized by the rapid development of single-cell DNA methylome assay technologies (27). In the past decade, most technologies have been based on pre-amplifying deaminated DNA (28-30) using random priming or single-strand adapter ligation before feeding amplified DNA to high-throughput sequencing. In addition, multiple enzymatic cytosine conversion methods have been developed to replace sodium bisulfite conversion to better preserve genomic DNA during library preparation (31). Inspired by these single-cell methods, we posit that similar preamplification methods and enzymatic conversion can also be used with Infinium arrays.

Besides the experimental challenges, current computational methods are not fully optimized for low-input DNA data. Current signal preprocessing practices may lead to low probe signal detection rates and probe over-masking. Signal detection has traditionally been determined by comparing probes’ signal intensities with negative control probes (32) or readings of Infinium-I out-of-band signals (33). A conservative threshold of the detection *p*-value was then determined to mask low-intensity probes. When the foreground and background signal intensities are separable, using a conservative threshold on high-input data would not harm detection sensitivity. This approach leads to a significant loss of true biological signals for limited DNA input. A method that maximally preserves biological signals while removing pure artifacts remains an unmet need.

Here, we systematically developed and evaluated experimental and computational methods to improve array sensitivity at low-input ranges and single cells. Our evaluation encompasses previously attempted adaptations, including using different bisulfite conversion elusions (34), using Formalin-Fixed Paraffin-Embedded (FFPE) restoration (35), combining DNA extraction (36) as well as methods that were never previously used with Infinium arrays, such as the enzymatic conversion and different preamplification strategies. We developed a new signal detection framework to address the computational challenge of processing data from limited DNA. We showed that this new method significantly improved array detection rates while effectively masking probes whose readings are dominated by background signals. We showed that the Infinium BeadChip is compatible with samples of low input down to single cells. And we presented end-to-end solutions to enable this technology for low-input and single-cell samples.

## MATERIALS AND METHODS

### Cell cultures

NIH3T3 (ATCC, CRL-1658) was obtained from American Type Culture Collection (ATCC) and cultured in DMEM (ATCC, 30-2002) containing 10% Calf Bovine Serum (ATCC, 30-2030) and 1% penicillin/streptomycin (Gibco, 15140122). B16-F0 (ATCC, CRL-6322) was obtained from ATCC and cultured in DMEM (ATCC, 30-2002) containing 10% Fetal Bovine Serum (Gibco, 45000-736) and 1% penicillin/streptomycin (Gibco, 15140122). All cells were maintained in a 37 °C incubator with 5% CO_2_ and cultured in a 75 cm^2^ culture flask (Fisher, BD353136).

### Cell flow sorting

5 × 10^6^ cell pellets of the NIH3T3 and the B16-F0 were resuspended in 50 μl of 0.1 µg/1 ml of 4,6-Diamidino-2-phenylindole (DAPI) (Sigma-Aldrich, D9542-5MG) in 1 ml of phosphate-buffered saline (PBS) (Life Technology, 10010023). Cells were filtered by a Falcon Cell Strainer Snap Cap (Falcon, 352235). DAPI-negative cells (1, 2, 5, 10, and 100 cells) from NIH3T3 and B16-F0 were sorted and collected into 96-well plates pre-loaded with 10 μl of 1X M-Digestion Buffer (Zymo Research, D5020-9) using a BD FACSAria^TM^ Fusion cell sorter (BD Biosciences) with a 100 μm nozzle.

### Mouse primordial germ cells

Gonads from embryonic Oct4-GFP transgenic mice (B6; B6;129S4-*Pou5f1^tm2Jae^*/J; Jackson Laboratory, strain #008214, RRID: IMSR_JAX:008214) were harvested at embryonic day E11.5, E12.5, E13.5, and E14.5. E0.5 was determined as the day of the copulatory plug detection (37). Gonads were dissected in calcium-and magnesium-free PBS (Gibco) and transferred into 500 μl of 0.25% Trypsin-EDTA (Gibco). Subsequently, the preparation of embryonic germ cells was carried out following the method previously described (38). For bisulfite mutagenesis, PGCs were snap-frozen for storage at -80°C until further processing.

### DNA extraction and bisulfite conversion

NIH3T3 and B16-F0 cells were harvested by centrifugation at 100g for 5 min at room temperature and washed twice using PBS (Gibco, 10010023). The DNeasy Blood and Tissue Kit (Qiagen, 69504) was used to extract genomic DNA from NIH3T3 and B16, according to the manufacturer’s protocol. DNA samples were quantified using Qubit 4.0 Fluorometer (Invitrogen) using the dsDNA HS Assay Kit (Invitrogen, Q33231). Bisulfite conversion was performed using three kits. DNA bisulfite conversion using EZ DNA Methylation Kit (Zymo Research, D5001) was performed according to the manufacturer’s instructions with the specified modifications for Illumina Infinium Methylation Assay. DNA bisulfite conversion using EZ DNA Methylation-Gold Kit (Zymo Research, D5005) and EZ DNA Methylation-Direct Kit (Zymo Research, D5020) was performed according to the manufacturer’s protocol. Cell lysis and bisulfite conversion from sorted cells and PGCs were performed with EZ DNA Methylation-Direct Kit according to the manufacturer’s instructions.

### DNA restoration

After bisulfite conversion, the bisulfite-converted DNA was eluted, resulting in an 8 µl volume. The Infinium HD FFPE DNA restoration kit (Illumina, WG-321-1002) was then used according to the manufacturer’s instructions. Following a 1-min incubation, the elution of the DNA was carried out using autoclaved ultrapure water for a 10 µl elution volume. The eluted DNA was stored at -20 °C before undergoing Infinium array processing. Intermediate DNA purifications were performed using the DNA Clean and Concentrator-25 Kit (Zymo Research, D4064).

### Cytosine conversion elusion size optimization

The Illumina Infinium Mouse Methylation BeadChip assays were conducted according to the manufacturer’s specifications with slight modifications (Supplemental Tables S1A and S1B). The original protocol specifies using 4 µl obtained from 12 – 22 µl eluted BCD, mixed with 4 µl 0.1 N sodium hydroxide for amplification and BeadChip reaction (Infinium HD Assay Methylation Protocol Guide 15019519 v01). However, commercial bisulfite conversion kits typically produce over 10 µl elusion in purifying the converted DNA, leading to only part of the eluted DNA (4 µl) used for the BeadChip assay. Previous studies have adjusted the elusion size or additional concentration steps to minimize DNA loss. For example, one option is to mix 7 µl eluted DNA with 1 µl 0.4 N sodium hydroxide (39-41). DNA input can be maximized by increasing NaOH concentration in denature step of the Infinium array. To preserve more input DNA, we compared four alternative combinations of elution buffer, input DNA volume, and NaOH concentration and volume.

### EM-array sample preparation

Libraries were prepared using the NEBNext Enzymatic Methyl-seq (NEB, E7120S) kit, following the manufacturer’s instructions. 50, 5, 2, or 0.5 ng of 5mC adaptor-ligated NIH3T3 DNA was used as input. HiFi HotStart Uracil+ Ready Mix (KAPA Biosystems, KK2801) was used to amplify the libraries following conversion before purification over SPRI beads (0.8x left-sided) and elution in nuclease-free water to yield final libraries. Libraries were then quantified by Qubit HS (Invitrogen, Q32851) and quality checked on an Agilent Bioanalyzer 2100 before sequencing on an Illumina MiSeq instrument to confirm conversion efficiencies. Details of each input’s elution size and library amplification cycles are listed in Supplemental Table S1C.

### ELBAR detection *p*-value calculation

In the standard Infinium BeadChip usage, >200 ng DNA is profiled, and probe detection calling is employed to filter out probes whose signals are subject to substantial background influence. However, this practice will cause significant biological signal loss from low-input datasets where users often seek to retain the most biological signal but can tolerate some background influence. To meet this need, we developed ELBAR (Eliminating BAckground-dominated Reading) to exclude/mask only probes lacking biological signals and entirely dominated by background signals. The ELBAR method is based on the observation that the beta value ranges depend on the probes’ total signal intensities and includes the following steps. First, we define total signal intensities as the sum of signals methylated (M) and unmethylated (U) alleles. Pooling in-band and out-of-band signals, ELBAR bins probes by log2-transformed M+U signal intensities. Next, ELBAR calculates the envelope of the beta value distribution for probes within each bin. The upper and lower bounds (calculated using the 5% and 95% quantiles to accommodate outliers) were traced to contrast the increase of M+U. Third, we define the background signal by looking for the first bin that deviates in the beta value envelope of the bin from the smallest M+U. Lastly, these probes’ maximum M and U signals were treated as the true background signal to compute detection *p*-values.

### Public datasets

The mouse MM285 datasets were downloaded from GEO under GSE184410 (42) (16). GEO accessions of other EPIC and HM450 datasets were provided in Supplemental Table S2A. BS-seq dataset for 4 PGC samples (E10.5, E11.5, one male E12.5, and one female E12.5) was downloaded from GEO under GSE76971 (43). TCGA testicular seminoma datasets (44) were downloaded from Genomic Data Commons (45).

### Infinium BeadChip data preprocessing and analysis

The Illumina Infinium BeadChips technology is based on sodium bisulfite conversion of DNA, with single base resolution genotyping of targeted CpG sites via probes on a microarray. Probes are designed to match specific 50 base regions of bisulfite-converted genomic DNA, with a CpG site at the probe’s 3’ end (46). Upon hybridizing with bisulfite-converted DNA, the probe undergoes a single-base extension that incorporates a fluorescently labeled ddNTP at the 3’ CpG site, enabling distinguishing of the C/T conversion resulting from bisulfite conversion. The fluorescent signals provide insight into the methylation status (methylated or unmethylated) of specific cytosine residues in the DNA sample. Signal intensity refers to the strength of the fluorescent signal emitted by the hybridized probes on the BeadChip. The signal intensity is directly related to the quantity of target molecules bound to specific probes. Probe success rates represent the proportion of successfully captured CpG probes for targeted CpGs.

The MM285 Infinium beadChip IDAT files underwent preprocessing, quality control, and analysis using the SeSAMe R package following preprocessing workflow (42). The probe detection *p-*value was computed using the pOOBAH algorithm, which leverages the fluorescence of out-of-band (OOB) probes. Subsequently, normalization was performed using noob, which applies a normal exponential deconvolution of fluorescent intensities based on the OOB probes. Additionally, a dye bias correction was applied using the dyeBiasNL function. Infinium Methylation BeadChip manifest and annotations data which include gene, chromatin state, sequence context, and other functional annotations, were obtained from http://zwdzwd.github.io/InfiniumAnnotation. Metagene plot was calculated using the KYCG_plotMeta function in the SeSAMe package. To calculate the F1 scores of each sample against the 250-ng control (205243950081_R01C01), a beta value greater than 0.5 was rounded to one and set to zero otherwise. Then, the F1 score is calculated by treating one as true and 0 as false and comparing the target sample with the 250- ng control, i.e., F1=2TP / (2TP+FN+FP) where TP is the true positive counts, FP is the false positive counts, and FN is the false negative counts.

## RESULTS

### Characterizing the suboptimal DNA signatures in public Infinium datasets

Suboptimal DNA quality and quantity impact Infinium methylation data through the manifestation of lower signal intensity and probe success rate. We first studied the probe success rate of over 100,000 public Infinium datasets deposited to GEO to identify the determinants of Infinium array performance (Fig 1A). Comparing DNA sources, we observed that bone, buccal, plasma cfDNA, esophagus, and saliva often yield data with suboptimal probe detection success. Further dataset stratification by sample preservation reveals that FFPE samples are significantly lower in probe success rate than non-FFPE samples. The lower signal intensity leads to skewing of beta values, which represent the methylation level at a specific site, towards an intermediate reading at 0.5 due to stronger relative influences of the signal background (Figs 1B – 1D) (33). This increase in the background signal influence is a continuous spectrum rather than a dichotomy of detection success vs. failure. When the input DNA is of high quality and quantity, most probes have stable and clustered signal intensities, leading to a bimodal beta value distribution. But in suboptimal datasets such as from FFPE and cfDNA samples, more probes carry lower signal intensities, leading to beta values approaching 0.5 (Fig 1C). This drop in sample quality in FFPE samples is likely intertwined with low DNA input amount, as supported by a similar transition of intermediate methylation readings in the low input data (Fig 1D and Supplemental Fig S1A). We also found that FFPE and cfDNA samples lose detection at different genomic regions. cfDNA, saliva, and buccal cells preferentially lose detection at GC-rich promoter sites, while FFPE samples are less biased across genomic territories (Fig 1E).

**Figure 1.**
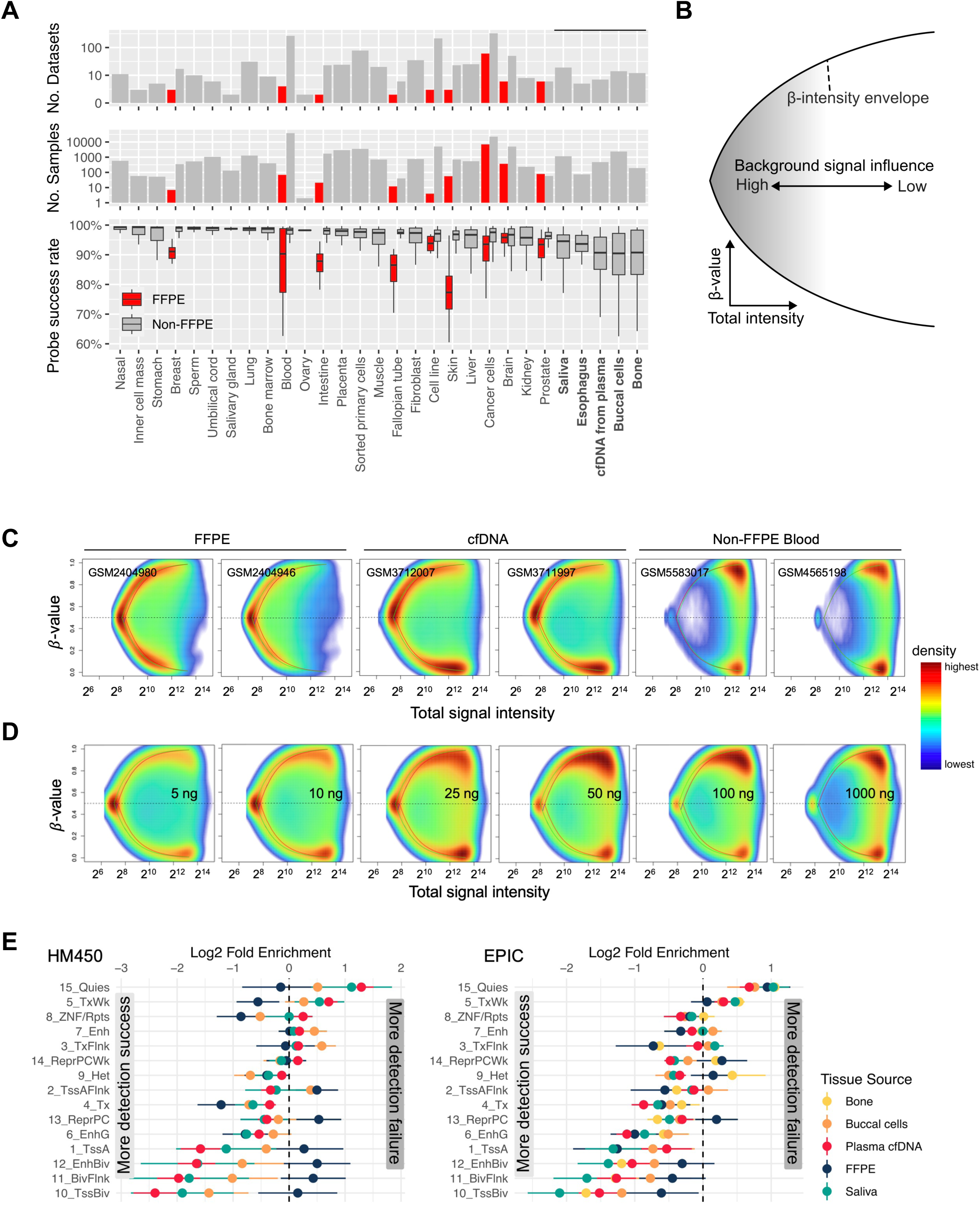
Lower input arrays exhibit lower signal intensity and lower probe success rates. (A) The probe success rate in 105,475 public Infinium methylation BeadChip array data sets. (B) Schematic showing the dependence of beta value on the probe’s total signal intensity. The x-axis represents the total signal intensity, and the y-axis represents beta values. (C) The intensity-beta plot compares different DNA sources and (D) input amounts in ng. (E) The probe detection rate of FFPE and cfDNA samples at different genomic regions from HM450 (left) array data and (right) EPIC array data.

### Infinium BeadChip workflows for low DNA input and single cells

To improve signal detection in low-input experiments, we developed 13 non-standard workflows using: 1) preamplification of the genomic DNA (see Supplemental Methods); and 2) enzymatic cytosine conversion methods, besides other protocol adjustments that preserve DNA load (see Fig 2A, Supplemental Tables S1A – S1C). We applied four workflows, including the standard (Workflow A) and three customized workflows, to mouse cell line DNA with input sizes ranging from 250 ng to single cells (Figs 2B – 2D, Supplemental Fig S2A). We found that the standard Illumina workflow can detect signals on 70% array probes with two ng DNA without modification, consistent with our prior characterization of the EPICv2 BeadChip (47). In the sub-2-ng range, the detection success rate drops rapidly for the unmodified workflow (Figs 2B, 2C, and S2B). Enzymatic base conversion (Workflow M) maximizes signal detection (84%) in 0.5 ng-input experiments, followed by a whole-genome preamplification-based method (Workflow J) and one with elution size modification alone (Workflow C). Due to the allelic nature of DNA, we use the F1 score (Methods), which binarizes beta values for comparison to the reference sample, rather than correlation. In the five-cell experiments, Workflow J (Figs 2B, 2D, and Supplemental Fig S2C) consistently reaches over 70% in signal detection (pOOBAH < 0.05) and close to 90% in the F1 score. Notably, Workflow J detects around 25% probes in single cells with an F1 score >70%, suggesting that the detected data is biologically informative despite a higher rate of detection failure. In contrast, the standard workflow is consistently under 50% in the F1 score, suggesting more biologically misleading readings.

**Figure 2.**
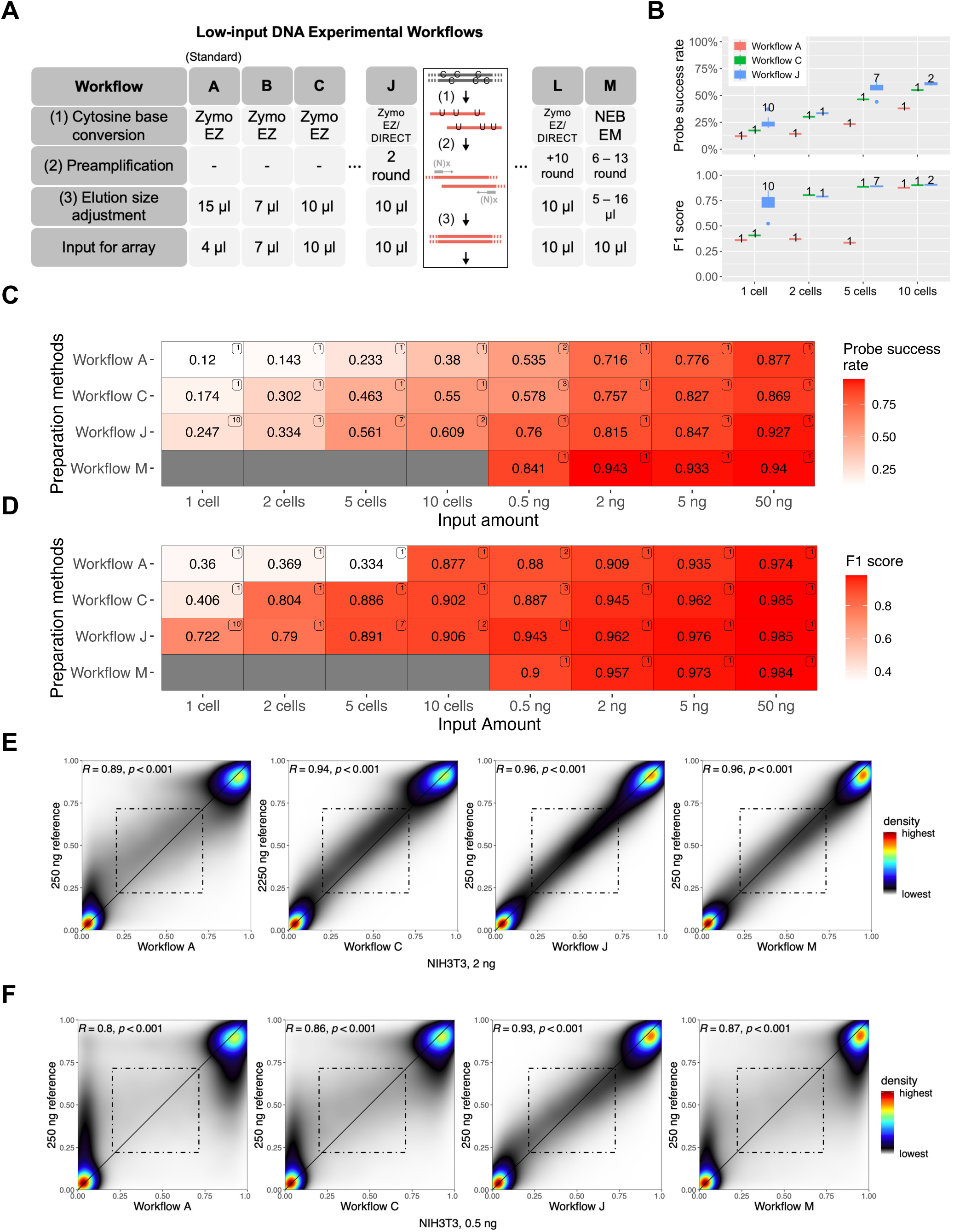
Infinium BeadChip performance in ultra-low input ranges. (A) A summary table of workflows used in this study. Workflow A is the Illumina standard workflow. (B) Box plots were used to visualize the probe success rates (top) and the F1 score (bottom). The number of samples for each experiment was displayed next to each box. (C – D) Comparison of four main preparation methods based on (C) probe success rate and (D) F1 score. The number on the top right corner of each tile indicates the number of samples analyzed in each experiment. See also Supplemental Fig S2. (E – F) Smooth scatterplots for the comparison of workflows A, C, J, and M with (E) 2 ng and (F) 0.5 ng of DNA input (*R*: spearman’s rho, p: p-value).

For 0.5 and 2 ng input ranges, Workflow J also improved the data quality as indicated by the higher correlation coefficient with a 250-ng dataset (Figs 2E, 2F, and Supplemental Fig S2D). The improvement is most evident in the recovery of intermediate DNA methylation, which is only observed in workflow J at 0.5 ng (Fig 2F), suggesting that preamplification helps all samples from fewer than ten cells to those with over 1 ng input (Fig 2E).

Previous studies have shown that FFPE restoration kits may improve signal detection on DNA of suboptimal quality. The FFPE restoration kit did not significantly improve the array performance with 50 ng and 0.5 ng non-FFPE DNA input (Supplemental Fig S2E). We also did not observe substantial performance differences among the three different bisulfite conversion kits (Supplemental Fig S2F). EZ-direct kit produced data of a slightly better probe detection rate likely due to the minimization of DNA loss from a single-step bisulfite conversion and purification.

We also compared preamplification based on random hexamers (N6) and random hexamers with a T7 primer at the 5’-end (N6-T7) for whole genome amplification (Supplemental Fig S2G) (48). The T7 sequence in N6-T7 serves as a second primer, allowing further PCR amplification. However, additional PCR cycles did not improve array performance (Supplemental Table S1D, Supplemental Figs S2B, and S2D). Moreover, the array with four Klenow amplification cycles (Workflow H) did not outperform the array with two cycles. Multiple amplifications were not found to yield better array performance. Given our findings, we followed the N6 amplification strategy for the subsequent analysis.

### Optimized workflow resolves intercellular heterogeneity while maintaining cell line identity

DNA methylomes profiled from a small number of cells often reveal cell-to-cell heterogeneity. We next tested whether our low-input method can uncover this heterogeneity and whether the cell population averages reduce to measurements from high DNA input. We merged the single and five-cell methylomes, respectively, and compared the combined data with the 250-ng methylome (Fig 3A). We found that the merged methylome reinstated the intermediate methylation measurements which are otherwise missing from the single low-input experiment representatives. Focusing on CpGs showing intermediate methylation in the 250-ng data, we observed a gradual dichotomization of methylation levels approaching 0% and 100% in the 10-cell data and more in the five-cell datasets (Fig 3B). This polarization is likely due to a higher genomic DNA homogeneity from reduced cell numbers. Given the same input cell number, preamplification (Workflow J) retained more intermediate readings (Fig 3C). The standard workflow (Workflow A) became nearly fully dichotomized at ten cells and struggled globally in the 5-cell experiment (Fig 3C).

**Figure 3.**
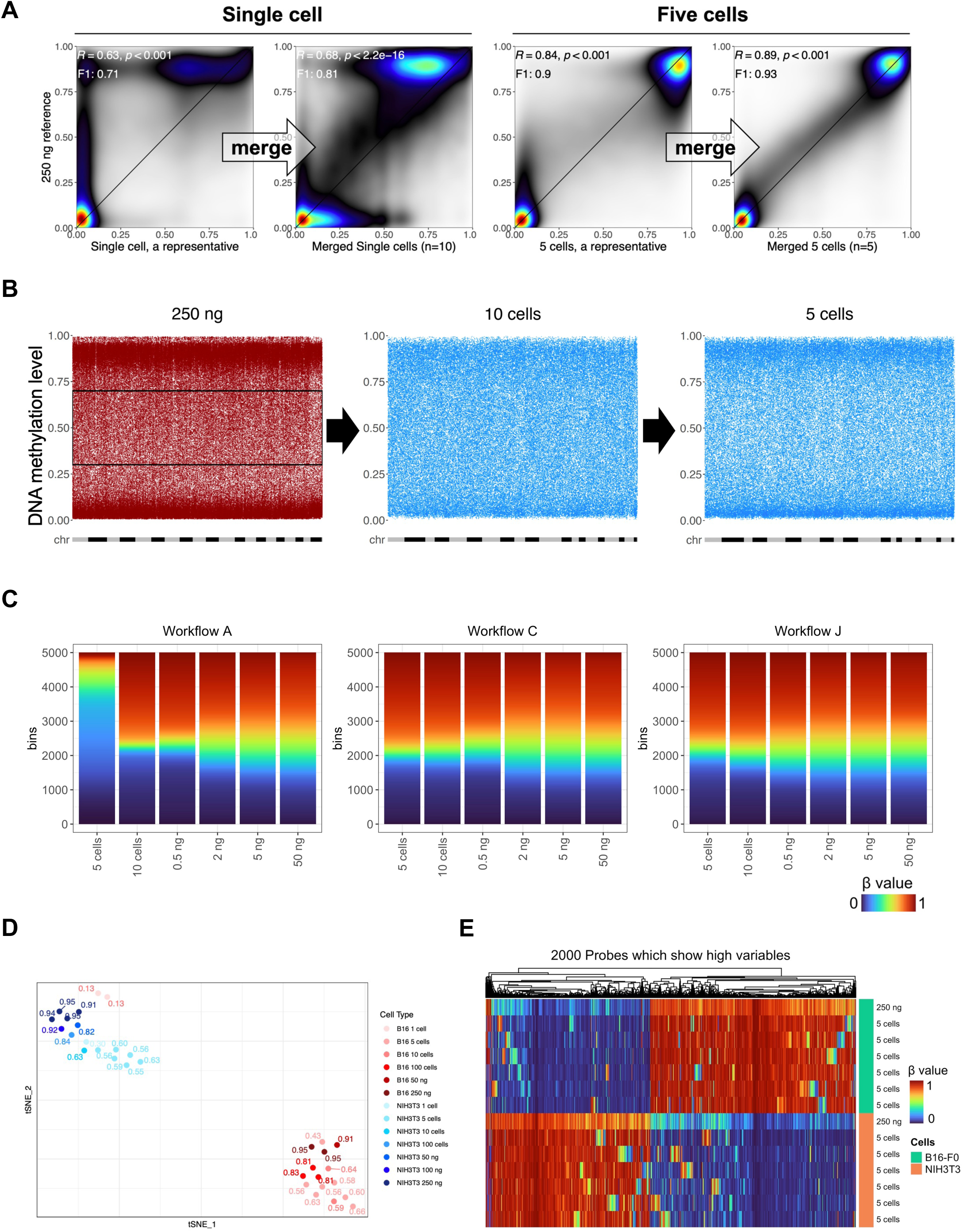
Low input methylation array data uncovered sample-to-sample heterogeneity. (A) Smooth scatter plots for methylomes from a representative single-cell dataset, the merged single-cell dataset (N=10), a representative 5-cell dataset, and the merged five-cell dataset (N=5), respectively, against the 250-ng input. (B) Distribution of intermediate methylation (0.3 to 0.7) of CpGs in the 250-ng methylome in 10- and 5-cell datasets. The X-axes were arranged in order from chromosome 1 to chromosome Y, with each chromosome displayed in alternating gray and black colors. (C) DNA methylation level distribution of 5 cells to 50 ng with different workflows. (D) tSNE cluster of NIH3T3 and B16 using 2,000 probes most variable in the methylation level. Fractions labeled in the plot are probe success rates for each dataset. All B16-F0, NIH3T3, and PGCs except 50 ng, 100 ng, and 250 ng were whole genomes amplified by Workflow I or J. (E) Heatmap showing array performance comparison of 5-cell methylomes of B16-F0 and NIH3T3 with 250-ng methylomes.

Despite the resolution of intermediate methylation levels, small-cell-number data largely retain global and focal methylation differences intrinsic to the cell type identity (Supplemental Fig S3A). All datasets from five to ten cells cluster with the 250 ng datasets of the same cell line on a t-distributed stochastic neighbor embedding (tSNE) projection (Fig 3D). Most single-cell samples are also grouped accordingly despite the erroneous placement of two single-cell B16 samples, likely due to an array-wide failure. The five-cell datasets are clearly separated in a metagene plot that suggested a higher global methylation for the B16 cells at all input ranges (Supplemental Fig S3B). Finally, the differentially methylated CpGs between the two cell lines are largely preserved in five-cell methylomes. Random discrepant methylation did occur more frequently at CpGs intermediately methylated in the 250 ng datasets for the two cell lines, respectively (Fig 3E), enriching for bivalent chromatin (Supplemental Figs S3C and S3D). Collectively, these data suggest the Infinium arrays can robustly profile five-cell samples. While the Infinium arrays can profile single cells, their performances are unstable (Fig 3D).

### GC-rich and high copy number regions retain detection in low input datasets

We next explored which genomic regions are most susceptible to detection loss in high and low input experiments. We first observed that signal intensities intrinsically depend on the probe sequences. We studied the within-sample intensity Z-score of HM450 autosomal probes across 749 normal samples from the TCGA cohort (Methods) (Fig 4A, Supplemental Fig S4A). Each violin plots all Z-scores of a probe in the cohort, and each data point represents a sample. The Z-score distribution for different probes shows clear probe dependence. Probes at the high and low signal intensity extremes have little overlap, suggesting a strong sequence dependence. Probes targeting GC-rich regions (as indicated by the number of “C”s in the probe sequences since “G”s are replaced by “A”s to pair with “T”s from bisulfite conversion) are associated with higher signal intensities (Fig 4B). This is supported by an enrichment of the detected probes in CpG islands (Figs 4C, 4D, and Supplemental Fig S4B), gene promoters, transcription factor binding sites, e.g., TFAP2C, and promoter-associated histone modifications, e.g., H3K64ac (Supplemental Figs S4C and S4D).

**Figure 4.**
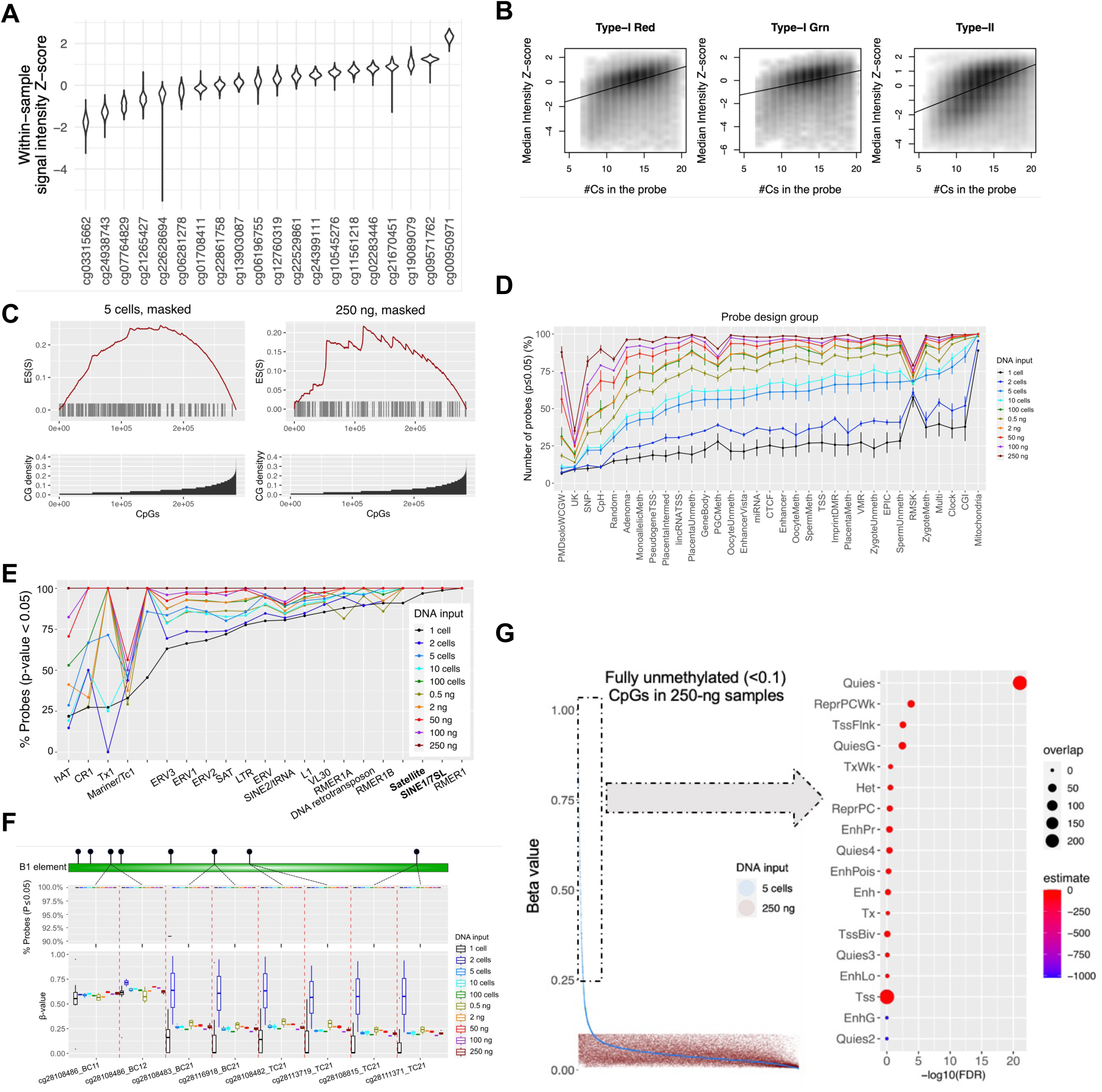
Low-input Infinium array preferentially loses detection on GC-sparse genome but retains detection on mitochondria and transposable elements. (A) Violin plot for intensity Z-score of representative HM450 autosomal probes across 749 normal samples from the TCGA cohort. (B) Correlation of signal intensities with the number of Cs in the probe sequence. (C) The enrichment of masked probes for low CpG density in NIH3T3 5-cell and 250-ng datasets. The x-axes represent the number of probes (N=297,415), arranged in ascending order of CpG density, and the y-axes represent the enrichment score. (D) The mean and standard deviation (with error bars) of the probe success rate of probe design groups (a complete description of the definition of each probe design group is listed in the legend of supplementary Fig S4) and (E) the mean of the probe success rate of transposable elements for varying DNA input amounts, ranging from single-cell to 250 ng datasets. The x-axes are organized in ascending sequence according to the probe success rates for five cells in (D) and single cells in (E). (F) (top) The probe detection rates of eight CpG probes targeting B1 elements ranged from single cells to 250 ng, and (bottom) corresponding beta values cover the same range. (G) (left) Beta values of the 250 ng and the five cells of NIH3T3; The x-axis is arranged in descending order of beta values of the five cells. (right) A dot plot for enriched ChromHMM (69) for the CpGs within the black squares (with a delta beta greater than 0.25 between the two samples) of the left figure.

Consistently, the probes that fail detection in 250-ng and 5-cell samples are significantly enriched in low-CpG density regions (Fig 4C). PMD solo-WCGWs---CpGs flanked by A/Ts and with no other CpGs within a 70-bp neighborhood at partially methylated domains (49)---are observed to lose most signal detection, consistent with their CpG-sparse nature. Probes targeting non-CG cytosine methylation also tend to lose detection in low-cell-number samples. Interestingly, mitochondrial CpG probes, transposable element CpGs, and other multi-mapping probes have the least probe detection loss in low-input datasets. The mitochondrial genome showed nearly 100% probe detection success in single, and two-cell experiments. This is likely due to the high copy number of mitochondrial genomes per cell (50). Similarly, other high copy number repetitive elements, such as the Satellite, B1 elements, and other SINE1/7SL elements, also show high probe success rates (Figs 4E and 4F). These results suggest that the multi-mapping probes may be used as a transposable element profiling tool for low-input samples.

### ELBAR preserves more signal detection for low-input datasets

The conventional detection *p*-value calculation aims at preventing false discovery in high-input datasets, where probes with suboptimal signal intensity are rare and a relatively clear decision boundary can be found. In low-input samples, more probes carry lower signal intensity and can overlap with measurements purely dominated by background signals (Supplemental Fig S5A). Applying the same detection *p*-value threshold may lead to a significant loss of biologically useful readings. To better balance sensitivity for low-input datasets, we developed the ELBAR algorithm, based on the observation that probes dominated by signal-background-only are always associated with intermediate methylation readings (Fig 5A). In brief, ELBAR looks for low-signal probes with intermediate methylation to model the background signal (Methods). Doing so can effectively remove background-induced artificial readings while minimally removing probes with biological signals (Fig 5B). In the cell line experiments, probes that survive ELBAR masking maintain a bimodal distribution of beta values as biologically expected. In contrast, pOOBAH, a prior method designed for high-input datasets, masked probes more aggressively. The probes surviving pOOBAH masking are slightly asymmetric in the beta-value envelope and show a small amount of background-dominated beta values (Fig 5B).

**Figure 5.**
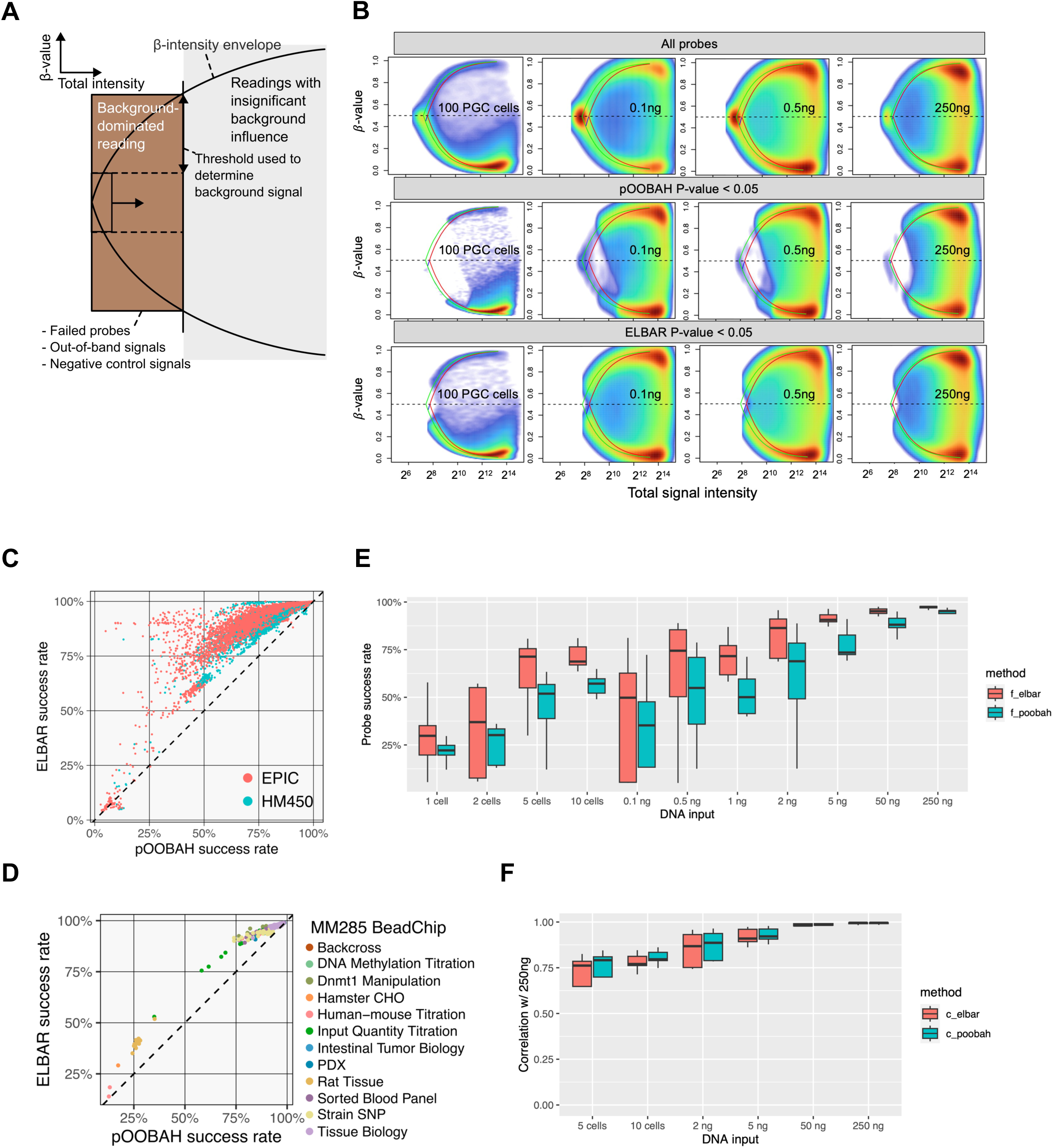
ELBAR detection preserves more probes with biological signals. (A) A schematic illustration of the ELBAR algorithm. The x-axis represents the total signal intensity, and the y-axis represents beta values. (B) Performance of ELBAR in eliminating background-dominated reading probes compared to pOOBAH and unfiltered in PGCs, 0.1-ng-, 0.5-ng-and 250-ng-input datasets. (C-D) ELBAR performance regarding the probe success rates for public human (C) and mouse (D) array datasets. (E-F) Comparison of ELBAR performance with pOOBAH regarding probe success rate (E) and Spearman’s rho (F) in low-input datasets with DNA input ranging from single cell to 250 ng.

Testing ELBAR on public EPIC, HM450 (Fig 5C, Supplemental Table S2A), and MM285 (Fig 5D) datasets, we found that it could rescue a significant number of probes compared to pOOBAH. Interestingly, experiments with array-wide failure remain low in detection rates, suggesting ELBAR can discriminate probe failure against array-wide failures. The probes rescued by ELBAR from pOOBAH show biological relevance, evidenced by higher correlation with the 250 ng datasets (Supplemental Fig S5B). Of note, ELBAR combines negative control probes for background calibration and only considers intermediate methylation reading from low-intensity probes. Hence, its masking would not be influenced by true biological methylations. For example, we validated ELBAR’s performance in samples with globally high, intermediate, and low methylation levels (Supplemental Figs S5C – S5E), including testicular seminoma tissues (Supplemental Fig S5F). ELBAR improves detection in wide input ranges (Fig 5E) without harming accuracy (Fig 5F).

### Low-input BeadChip data captures the demethylation dynamics in primordial germ cells

PGCs are typically present in low numbers, hindering their analysis by the standard Infinium array processing workflow (51-53). In mammals, PGCs undergo genome-wide epigenetic reprogramming, including global DNA methylation loss, as they migrate from the epiblast to the bipotential gonads (54). This corresponds to embryonic day(E)7.5 to E14.5 of development in the mouse. We applied our optimized method to study the methylation of mouse gonadal PGCs collected at E11.5 to E14.5 (Fig 6A). For each time point, PGCs from a pair of embryonic gonads were FACS sorted (Methods), and the aliquoted volume varied from 0.25 µl to 9 µl. We employed workflow J, with pre-amplified DNA amounts ranging from ∼1 ng to 13 ng (Supplemental Table S2B). As a contrast, we included methylome profiles of mouse liver, lung, ovary, and testes tissues in our analysis (Methods, Supplemental Table S2C).

**Figure 6.**
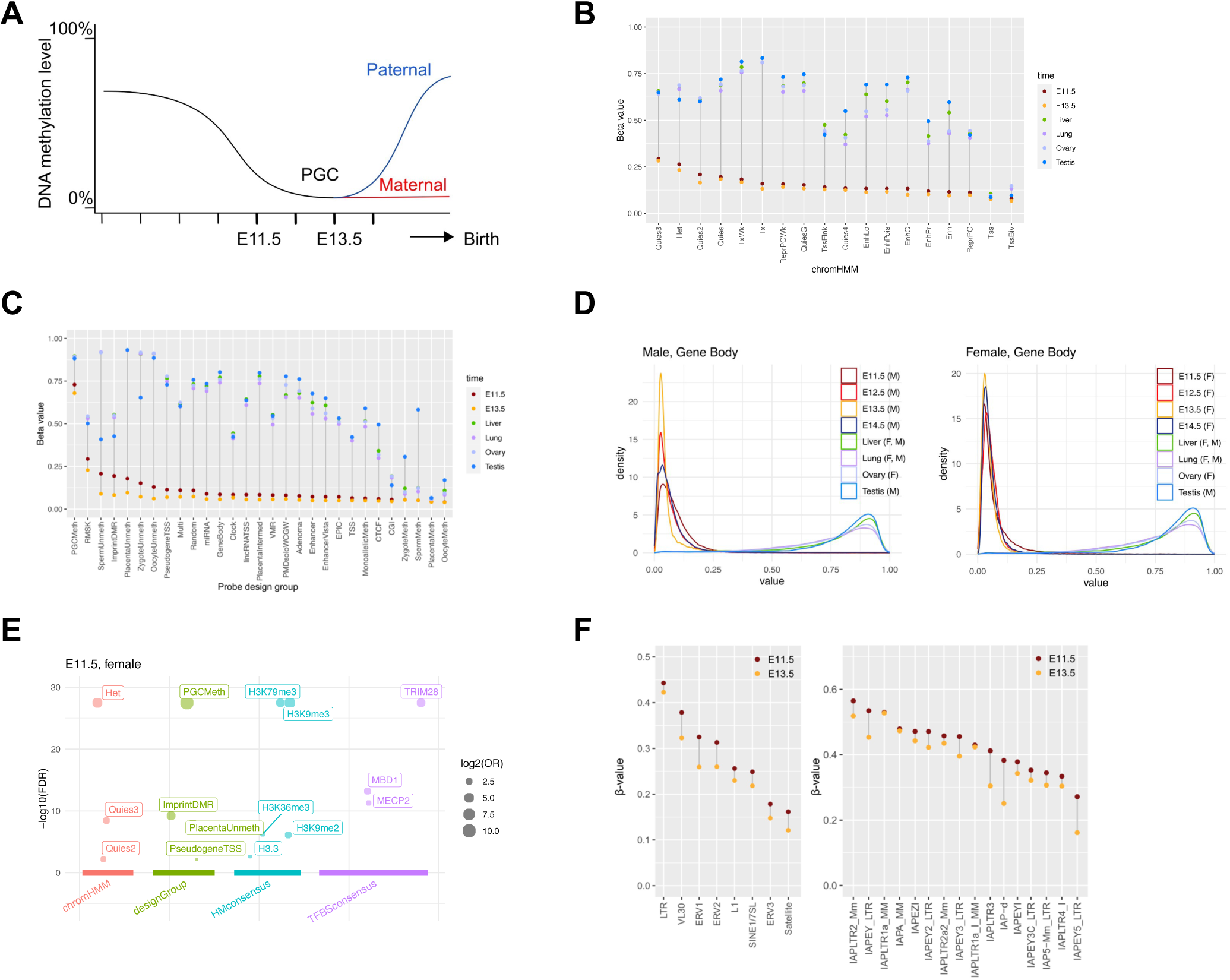
Low-input Infinium array data reveals epigenetic dynamics of primordial germ cell development. (A) Expected DNA methylation dynamics during primordial germ cell developments. E11.5 and E13.5 refer to embryonic days after fertilization. (B-C) DNA demethylation patterns in PGCs and tissues stratified by ChromHMM states (B) and mouse array probe design groups (C). (D) DNA methylation distribution in gene body regions in PGCs and tissues. M: male; F: female. (E) Enriched genomic features by CpGs that retain methylation in the three female E11.5 PGCs. Query CpGs are probes with beta values over 0.5 in the three samples. (F) DNA demethylation patterns in transposable elements (left) and IAP elements (right) across PGCs and tissues.

Consistent with prior knowledge, PGCs exhibit the lowest methylation level relative to somatic tissues and adult gonads across major chromatin states (Figs 6B and 6C). Consistent with the probe design rationale, “PGCMeth” probes showed resistance to methylation in E11.5 and E13.5 PGCs. The genome-wide distribution of PGC methylation loss is largely uncoupled from their methylation states observed in non-PGC tissues. For example, the genomic regions with active gene transcription (ChromHMM states “Tx” and “TxWk”) were associated with the highest methylation in non-PGC tissues (Fig 6C). But in PGCs, gene bodies are less methylated than heterochromatin and transcriptionally quiescent regions (Figs 6B and 6C). We observed a partial methylation loss in E11.5 PGCs. The demethylation process culminated in E13.5 PGCs. Male PGCs initiated re-methylation as early as E14.5, while female PGCs remained similarly unmethylated at E14.5 as E13.5. This is consistent with *de novo* methylation occurring in female germline postnatally as oocytes are recruited for growth during each reproductive cycle (55). This sex disparity in methylation rebound is evident in imprinted gene-associated differentially methylated regions (DMRs) and gene bodies (Fig 6D and Supplemental Fig S6A).

The arrays allow for detailed analysis of the timing of the DNA methylation change across genomic elements. For example, the retained DNA methylation at heterochromatin is enriched at TRIM28 binding sites (Fig 6E and Supplemental Fig S6B). TRIM28 regulates transcription of transposable elements, particularly endogenous retroviruses (ERV) (56). Specific ERV elements are known to retain DNA methylation in human PGC development (57) and mice (58). These observations suggest a critical role for DNA methylation in TE suppression for maintaining germ cell genome integrity and intergenerational epigenetic inheritance.

Interestingly, PGC residual methylation is also enriched for the binding of the DNA methylation reader proteins MECP2 and MBD1, supporting their reported roles in DNA methylation-mediated TE regulation (59). Among TE DNA families, LTR elements were more resistant to PGC demethylation than SINEs and LINEs, consistent with their functions as transcriptional promoters (60). The role of methylation retention in TE regulation is further supported by the higher methylation retention in evolutionarily younger TE subfamilies than older TE families (61) (Fig 6F). IAP elements were previously highlighted to exhibit extensive methylation retention in PGCs (62). We identified a heterogeneous pattern of their demethylation dynamics in PGCs. IAPLTR2, IAPEY, and IAPA are among the most resistant families, while IAPEY5, IAPLTR4, and IAP5 are the least methylated.

In addition to transposable elements, imprinted gene DMRs, germ cells, and placenta-specific hypomethylated sites show later DNA methylation loss compared to the rest of the genome. This is due to active DNA demethylation pathways, mediated by TET proteins, that are required for methylation erasure at imprinting control regions (63,64). CGs flanked by A/Ts are more susceptible to aging-associated DNA demethylation (49). We did not observe this sequence context preference for PGC development (Supplemental Fig S6C), suggesting a distinct, TET-mediated demethylation mechanism.

## DISCUSSION

Despite the success of Infinium BeadChips in DNA methylome profiling, their potential for “difficult” DNA, i.e., when the input is limited in quality, quantity, or both, has not been fully explored or optimized. This restricts the scope of the Infinium array usage for cell-free DNA, microdissected tissues, and other samples of limited availability. Here, we presented experimental and computational resources to enable array usage in these suboptimal settings, especially with low-input DNA. Experimentally, we explored the array’s compatibility with random priming-based whole genome preamplification and enzymatic base conversion by TET/APOBEC3A. Our data suggests that both preamplification and enzyme-based conversion significantly improved the array data quality. Computationally, we developed ELBAR for preserving biological signals from suboptimal input datasets. ELBAR excludes only probes dominated by background noises.

Besides, we comprehensively characterized the biological and technical determinants of array performance from public datasets. From surveying 100,000+ datasets and using probe detection rate as the main performance metric, we found that cell-free DNA, saliva, bone, and FFPE samples, tend to have worse detection rates compared to cultured cells, primary and fresh frozen tissues. FFPE samples and those of sheer lower input show a different genomic distribution in signal detection. Plasma cfDNA tends to cause detection loss at CpG-sparse, GC-low genomic regions and preserved detection at CpG-rich regions such as bivalent chromatin. In contrast, FFPE-induced detection loss is less biased across genomic regions. This low-input sample bias can be attributed to the weaker intrinsic signal from GC-low probes (Figs 4C and 4D). As cfDNA localization is known to be linked to nucleosome footprints and can inform cell of origin (65), the array signal intensity bias may serve as an unconventional source of cell of origin signal to complement the methylation signal that the array data already carries.

Before this work, the Infinium BeadChips are known to have reasonable low-input performance down to 10 ng (34). This is due to the isothermal amplification in the standard workflow. The lower input limit of the technology remained unexplored. It was also unclear to what extent precision and sensitivity would be compromised with decreased DNA input. Our work shows that Infinium arrays are reasonably compatible with picogram-range input DNA or single-digit cell number. Preamplification using Klenow fragments and enzymatic conversion further magnifies probe signal intensities, leading to reproducible profiling of five-cell methylomes. Interestingly, additional adapter-based PCR amplification did not lead to a further increase in probe detection or accuracy (Supplemental Table S1D), likely due to loss of library complexity from amplification bias (66).

The Infinium technology is currently not cost-competitive for single-cell DNA methylome profiling due to the incompatibility with sample multiplexing (67). Although the focus of this work is to explore whether these arrays can profile samples of limited quantity, such as microdissected tissues (23), rather than using them as single-cell profiling methods, preamplification is shared between sequencing-based methods using Klenow or other polymerases (26). Infinium BeadChip data reflects the allelic nature of the DNA methylation signal on low DNA input. Intermediate methylation levels were resolved to high and low methylation readings. These binary readings reflect cell-to-cell heterogeneity and when merged, their population averages recapitulated bulk input measurements. Our single-cell array data reached 20% detection, similar to previous deep single-cell WGBS datasets (29).

Previous data analysis workflows relied on a single threshold for determining signal detection success (68). While this is a viable assumption for the high input data, it does not always hold for low-input datasets. In low-input datasets, biological signals overlap more with background signals in signal intensity, particularly for probes with intrinsically low foreground signals. Our analysis showed that this intrinsic propensity is linked to the number of Cs in the probe sequences, likely reminiscent of a GC content bias as the probes are designed to be G-less to pair with converted genomic DNA. The stronger overlaps of biological with background signal not only obfuscate the detection discrimination but also bias the beta value calculation towards 0.5 due to the incomplete subtraction of signal background from the observed compound signal. Users should consider this major tradeoff of measurement precision for sensitivity.

Compared to pOOBAH and other detection calling methods designed to minimize false discovery in high-input datasets, ELBAR seeks to mask only probes fully dominated by signal background, leaving probes with true biological signals visible in downstream analysis. However, we caution that probe readings surviving ELBAR detection may be influenced by background signals to various degrees, leading to unstable quantitative accuracy. The detection *p*-values can serve as measures of background influences besides their traditional use for probe masking.

Despite the reduced probe detection in low-input datasets, probes that target multi-copy DNA, such as mitochondria and repetitive elements (e.g., the B1 elements and satellite sequences), retain high signal intensities. In the low-input datasets, we observed that these probes measure aggregated signals from multiple genomic loci, making an unconventional use of the methylation BeadChips---as a tool to study the global epigenetic regulation of multi-copy TEs. In our work, we applied our low-input protocol to profile mouse PGCs. We validated the dynamics of global methylation erasure in PGCs, a sex disparity in remethylation, as well as the demethylation resistance at TRIM28 binding sites which are known to escape germ cell epigenetic remodeling (57). These multi-mapping probes also revealed that evolutionarily younger LTR repeat families retained more methylation than other repeat families. These methylation retentions can protect germ-line genome integrity from TE transcriptional mobilization.

## CONCLUSION

We presented experimental and computational solutions for applying Infinium BeadChips to low-input and single-cell samples. Based on whole-genome preamplification and enzymatic base conversion, our new methods revealed a previously underappreciated low-input potential of this popular methylation profiling assay. We demonstrated the power of these methods by applying them to uncover detailed demethylation dynamics of murine primordial germ cell development.

## DATA AVAILABILITY

All BeadChip data produced in this study is available through GEO (accession: GSE239290). ELBAR and other informatics for low input methylation BeadChip are implemented in the SeSAMe (version 1.18.4) available through Bioconductor (https://doi.org/doi:10.18129/B9.bioc.sesame).

## SUPPLEMENTARY DATA

Supplementary Data are available at NAR online.

## ACKNOWLEDGEMENTS

We thank the Center for Applied Genomics Genotyping Core at the Children’s Hospital of Philadelphia for their help with array processing.

