## Supplementary Files for "Low-input and single-cell methods for Infinium DNA methylation BeadChips"

### SUPPLEMENTARY FIGURES

# S1

**A**

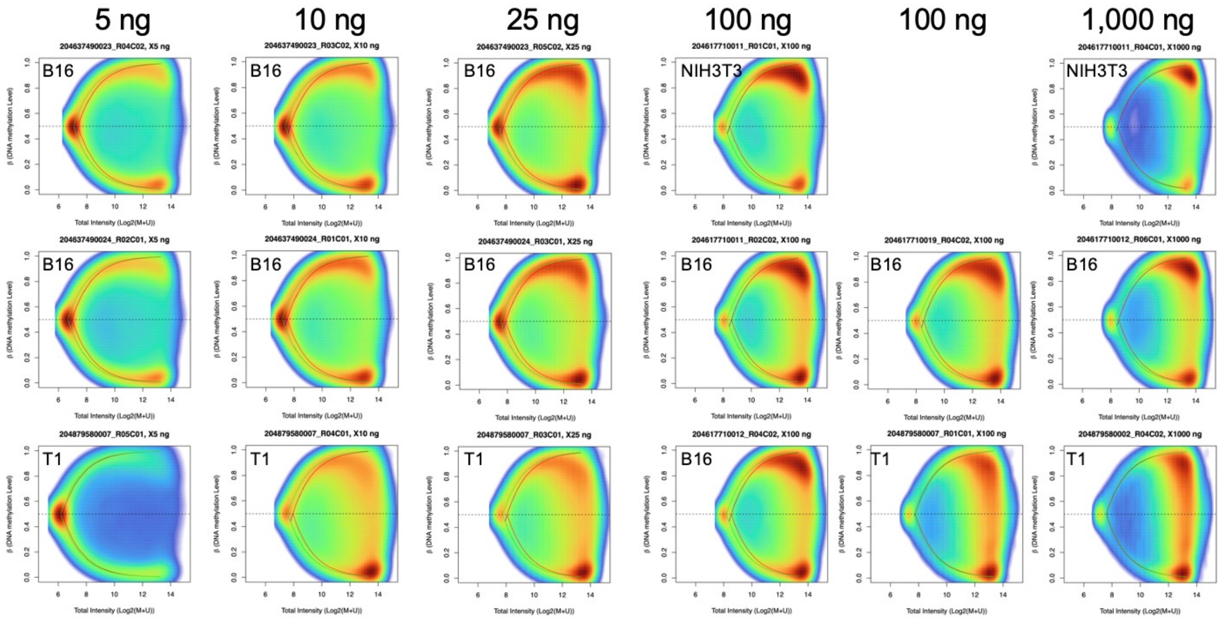

**Supplemental Figure S1.** (A) Signal intensity and beta values comparing among 5 ng to 1 µg of B16 (ATCC, CRL-6322), NHI3T3 (ATCC, CRL-1658), and T1.

T1: A small Intestine of a male mouse of 129S1_SvImJ

# S2

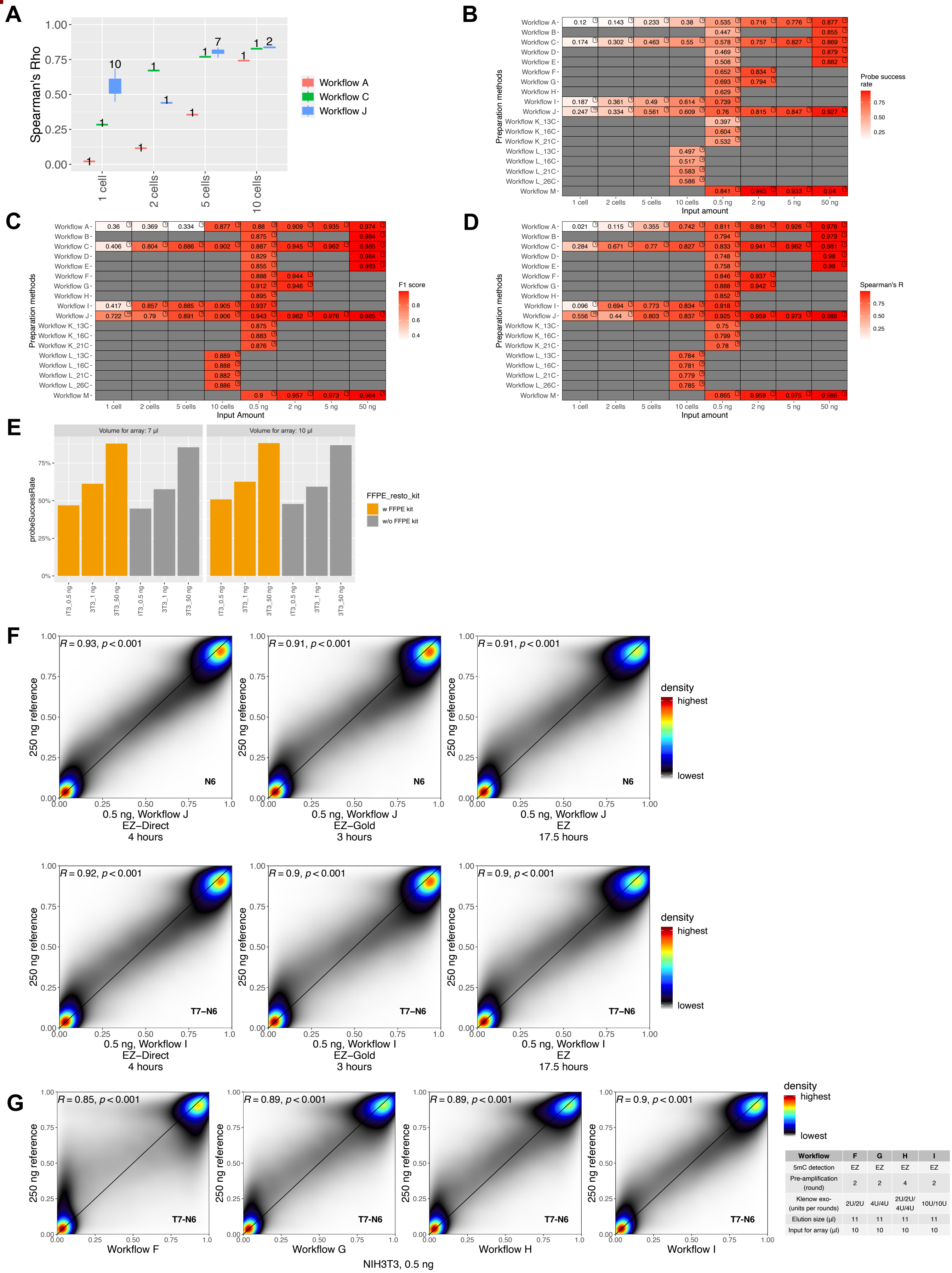


| Workflow | F | G | H | I |
|---|---|---|---|---|
| 5mC detection | EZ | EZ | EZ | EZ |
| Pre-amplification (round) | 2 | 2 | 4 | 2 |
| Klenow exo- (units per rounds) | 2U/2U | 4U/4U | 2U/2U/ 4U/4U | 10U/10U |
| Elution size (µl) | 11 | 11 | 11 | 11 |
| Input for array (µl) | 10 | 10 | 10 | 10 |

**Supplemental Figure S2.** (A) A box plot illustrates Spearman's Rho for three workflows among 1 to 10 cells. (B – D) Comparison of preparation methods based on (B) Probe success rate, (C) F1 scores, and (D) Spearman's rho. The number displayed on the top right of each tile in B – D indicates the total count of samples analyzed in each experiment. (E) FFPE restoration kit performance comparison. w FFPE kit: ATCC_NIH3T3 samples with treat FFPE kit (WG-321-1002); w/o FFPE kit: Experiment without FFPE kit treatment. (F) Bisulfite conversion kits comparison with workflows I and J. Bisulfite conversion kits and BCD preparation times were given in x-axes. 2 cycles of whole genome amplification and purification process for all these workflows were 3 hours. (G) Smooth scatterplots for comparing sample preparation methods with 0.5 ng of DNA input (*R*: spearman's rho, p: p-value). All experiments were performed with the EZ DNA methylation kit.

S3

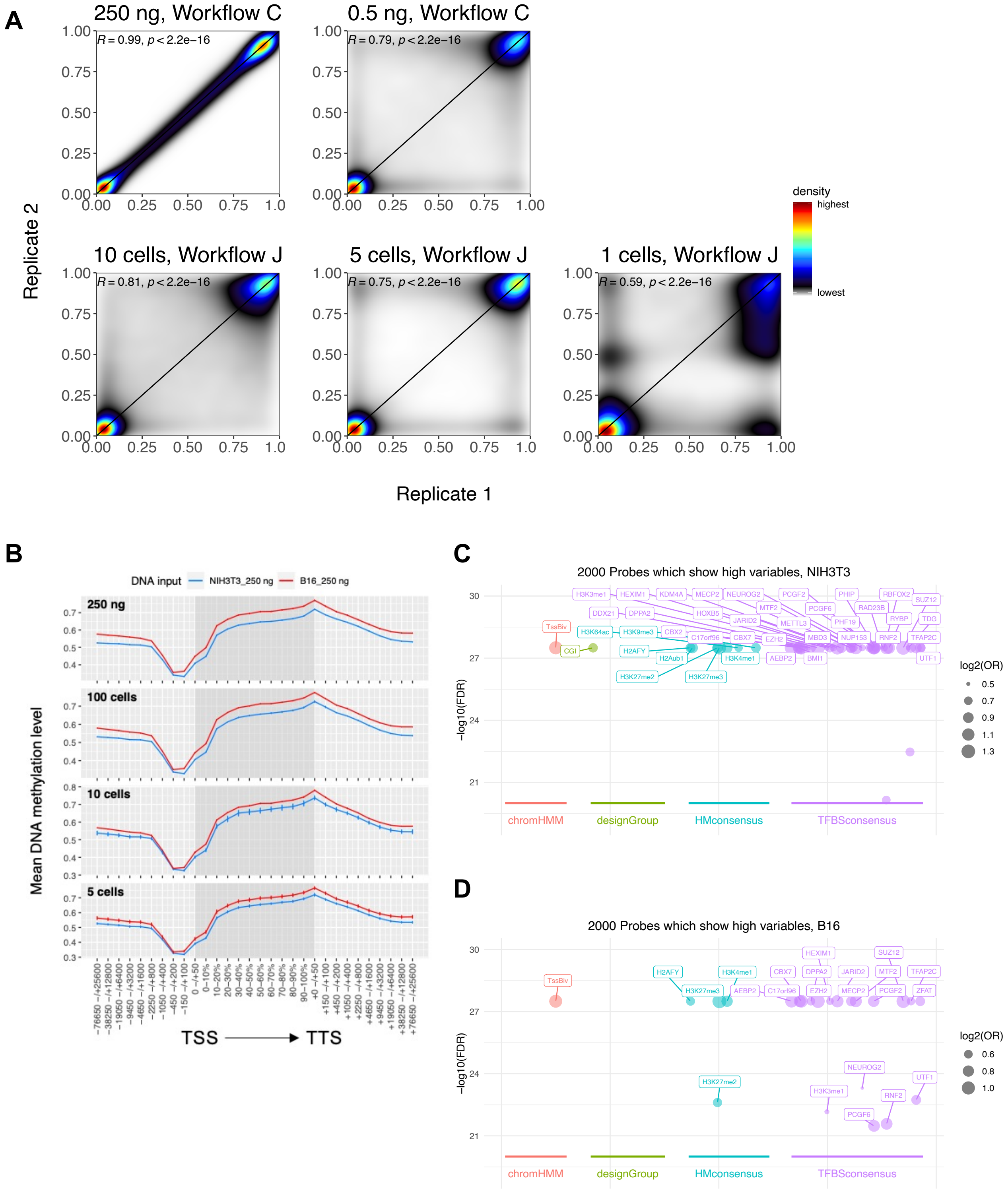
A
250 ng, Workflow C
0.5 ng, Workflow C
10 cells, Workflow J
5 cells, Workflow J
1 cells, Workflow J
R = 0.99, p < 2.2e−16
R = 0.79, p < 2.2e−16
R = 0.81, p < 2.2e−16
R = 0.75, p < 2.2e−16
R = 0.59, p < 2.2e−16
Replicate 2
Replicate 1
density
highest
lowest
B
DNA input
NIH3T3_250 ng
B16_250 ng
250 ng
100 cells
10 cells
5 cells
Mean DNA methylation level
TSS
TTS
C
2000 Probes which show high variables, NIH3T3
−log10(FDR)
log2(OR)
chromHMM
designGroup
HMconsensus
TFBSconsensus
D
2000 Probes which show high variables, B16
−log10(FDR)
log2(OR)
chromHMM
designGroup
HMconsensus
TFBSconsensus

**Supplemental Figure S3.** (A) Smooth scatterplots for comparison correlation among technical replicates. (B) The mean beta distribution of covered CpGs was determined around transcription start sites (TSS) using NIH3T3 and B16-F0. GRCm38 (mm10) reference genome was split with a length of 100 bp, and methylation levels were average methylation levels on each bin. (C – D) Enrichment of 2,000 highly variable CpGs by DNA methylation in 5 cells of (C) NIH3T3 and (D) B16-F0.

**S4**

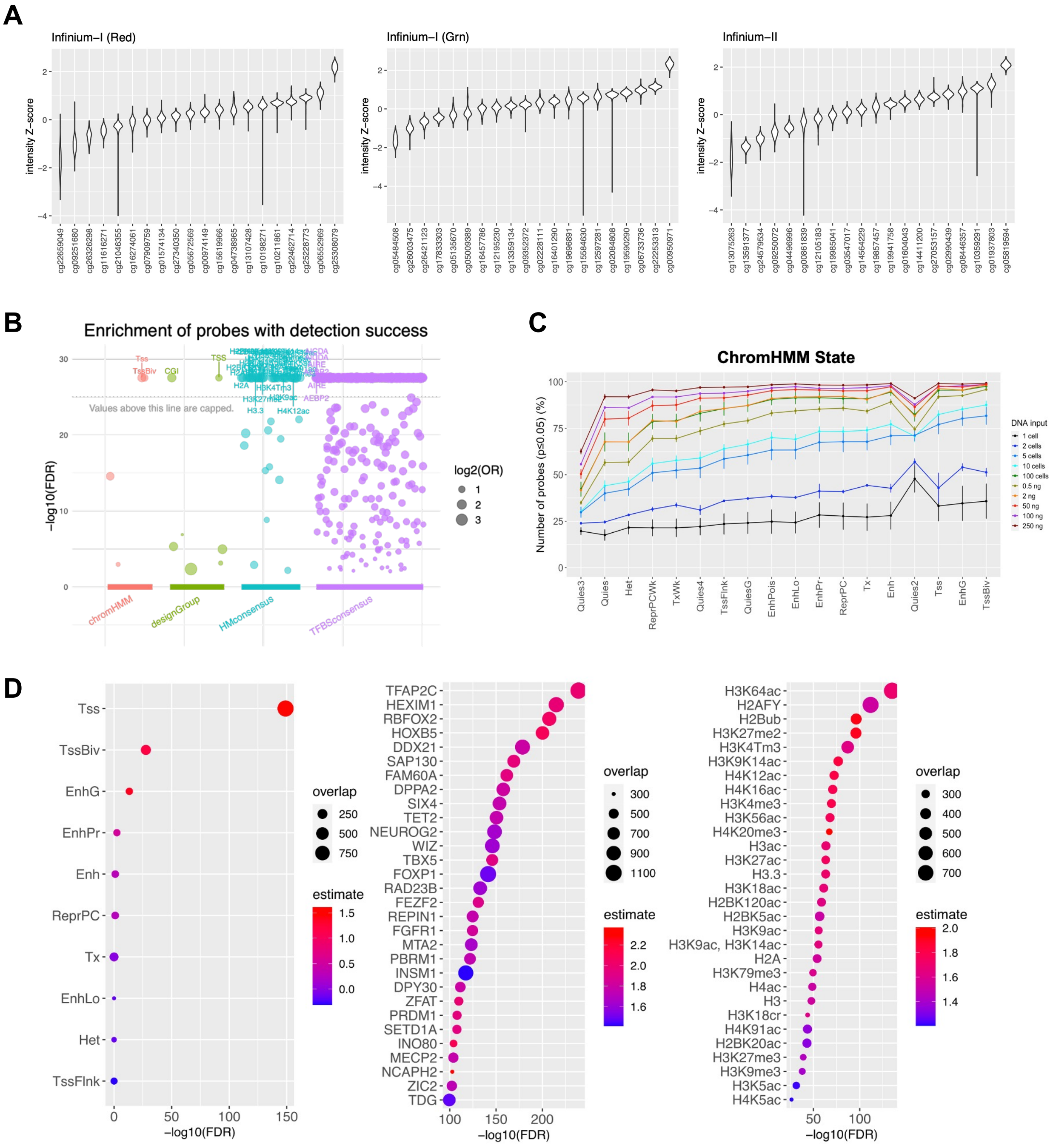

A
Infinium−I (Red)
Infinium−I (Grn)
Infinium−II
intensity Z−score
B
Enrichment of probes with detection success
Values above this line are capped.
−log10(FDR)
log2(OR)
chromHMM
designGroup
HMconsensus
TFBSconsensus
C
ChromHMM State
Number of probes (p≤0.05) (%)
DNA input
1 cell
2 cells
5 cells
10 cells
100 cells
0.5 ng
2 ng
50 ng
100 ng
250 ng
Quies3
Quies
Het
ReprPCWk
TxWk
Quies4
TssFlnk
QuiesG
EnhPois
EnhLo
EnhPr
ReprPC
Tx
Enh
Quies2
Tss
EnhG
TssBiv
D
Tss
TssBiv
EnhG
EnhPr
Enh
ReprPC
Tx
EnhLo
Het
TssFlnk
overlap
estimate
−log10(FDR)
TFAP2C
HEXIM1
RBFOX2
HOXB5
DDX21
SAP130
FAM60A
DPPA2
SIX4
TET2
NEUROG2
WIZ
TBX5
FOXP1
RAD23B
FEZF2
REPIN1
FGFR1
MTA2
PBRM1
INSM1
DPY30
ZFAT
PRDM1
SETD1A
INO80
MECP2
NCAPH2
ZIC2
TDG
H3K64ac
H2AFY
H2Bub
H3K27me2
H3K4Tm3
H3K9K14ac
H4K12ac
H4K16ac
H3K4me3
H3K56ac
H4K20me3
H3ac
H3K27ac
H3.3
H3K18ac
H2BK120ac
H2BK5ac
H3K9ac
H3K9ac, H3K14ac
H2A
H3K79me3
H4ac
H3
H3K18cr
H4K91ac
H2BK20ac
H3K27me3
H3K9me3
H3K5ac
H4K5ac

**Supplemental Figure S4.** (A) Violin plots for intensity Z-score of HM450 autosomal probes by assay types. (B) Enrichment analysis for CpGs probe in which detection succeeds. (C) The mean and standard deviation (with error bars) of the probe success rate by chromHMM status for varying DNA input amounts, ranging from single cells to 250 ng. X-axis is arranged in ascending order based on the five-cell probe success rates. (D) Dot plots of enrichment region for ChromHMM (left), transcription binding sites (middle), and histone modification (right) in not masked CpG probes were shown for the single cell to 250 ng input array of B16 and 3T3.

Adenoma: CpGs for colon cancer-specific methylation changes; CGI: CpGs in CpG islands; Clock: CpGs that age associated; CpH: probes for CpH sites; CTCF: CpG probes to capture the CTCF binding sites; Enhancer: CpGs to represent tissue-specific enhancer elements; EnhancerVista: CpGs to cover enhancers validated in the VISTA database (1); EPIC: human EPIC array CpGs; GeneBody: CpGs in gene body regions; ImprintDMR: CpGs for imprinted gene differentially methylated regions; LincRNATSS: CpGs for linc RNA TSS; miRNA: miRNA CpGs; Mitochondria: CpGs for mitochondrial DNA.; MonoallelicMeth: CpGs potentially involved in mono-allelic methylation; Multi: multi-mapping probes; OocyteMeth: CpGs observed to be specifically methylated of oocytes; OOcyteUnmeth: CpGs observed to be specifically unmethylated of oocytes; PGCMeth: CpGs observed to be specifically methylated of primordial germ cells; PlacentaIntermed: CpGs that displayed intermediate methylation level in a placenta tissue sample (2); PlacentaMeth: CpGs observed to be specifically methylated of placental tissues; PlacentaUnmeth: CpGs observed to be specifically unmethylated of placental tissues; PMDsoloWCGW: solo-WCGW at PMD CpGs; PseudogenTSS: CpGs in pseudogene transcription sites; RMSK: RepeatMasker probes; SNP: strain-specific SNPs: Random: randomly selected CpGs; SpermMeth: CpGs observed to be specifically methylated of Spermatocytes; SpermUnmeth: CpGs observed to be specifically unmethylated of Spermatocytes; TSS: CpGs for TSSs; UK: non-informative probes; VMR: CpGs for variably methylated regions; ZygoteMeth: CpGs observed to be specifically methylated of zygotes; ZygoteUnmeth: CpGs observed to be specifically unmethylated of zygotes.

S5

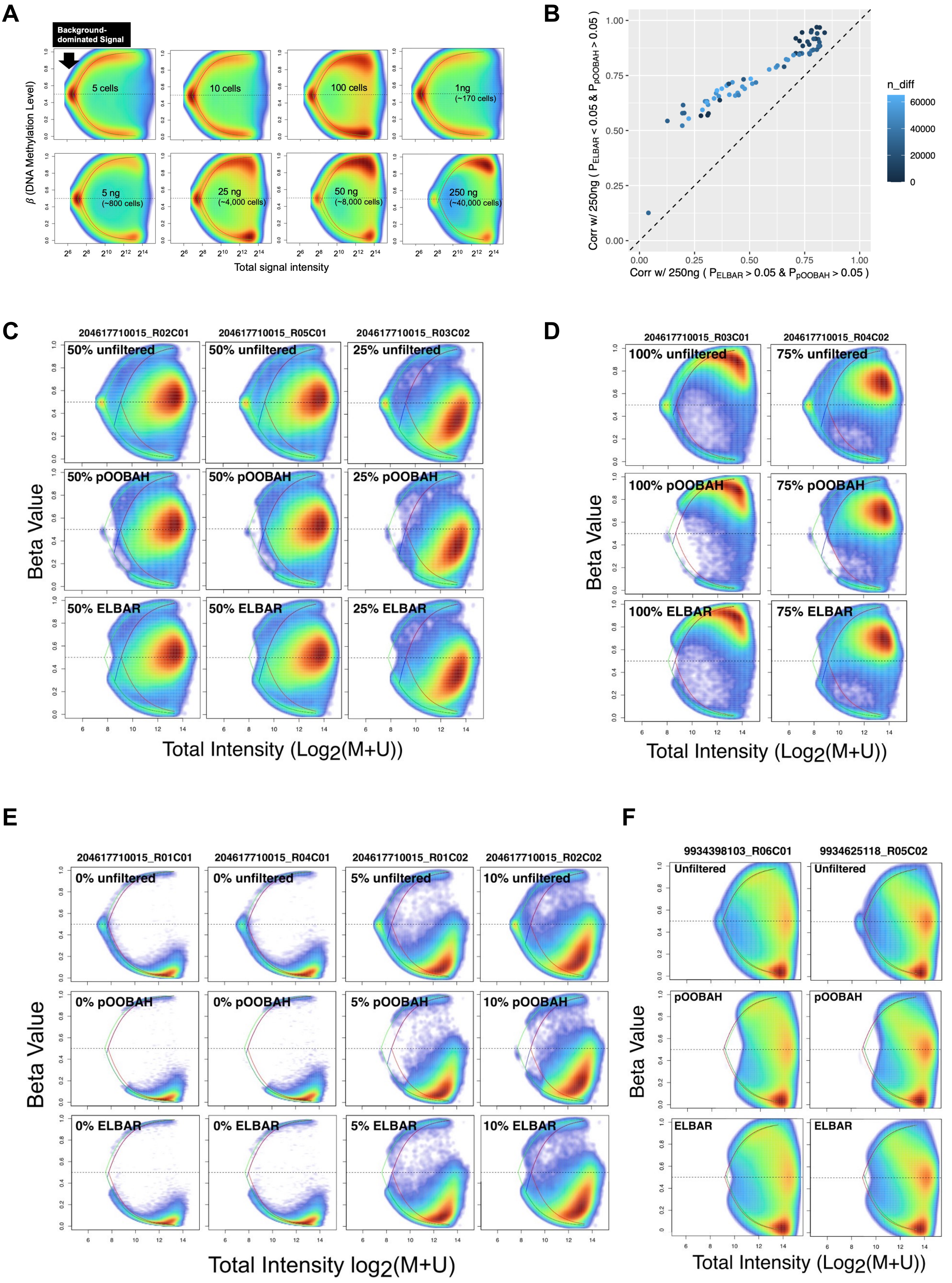
A
B
Background-dominated Signal
β (DNA Methylation Level)
5 cells
10 cells
100 cells
1ng (~170 cells)
5 ng (~800 cells)
25 ng (~4,000 cells)
50 ng (~8,000 cells)
250 ng (~40,000 cells)
Total signal intensity
Corr w/ 250ng ( PELBAR < 0.05 & PpOOBAH > 0.05 )
Corr w/ 250ng ( PELBAR > 0.05 & PpOOBAH > 0.05 )
n_diff
60000
40000
20000
0
C
204617710015_R02C01
204617710015_R05C01
204617710015_R03C02
50% unfiltered
25% unfiltered
50% pOOBAH
25% pOOBAH
50% ELBAR
25% ELBAR
Beta Value
Total Intensity (Log2(M+U))
D
204617710015_R03C01
204617710015_R04C02
100% unfiltered
75% unfiltered
100% pOOBAH
75% pOOBAH
100% ELBAR
75% ELBAR
E
204617710015_R01C01
204617710015_R04C01
204617710015_R01C02
204617710015_R02C02
0% unfiltered
5% unfiltered
10% unfiltered
0% pOOBAH
5% pOOBAH
10% pOOBAH
0% ELBAR
5% ELBAR
10% ELBAR
Total Intensity log2(M+U)
F
9934398103_R06C01
9934625118_R05C02
Unfiltered
pOOBAH
ELBAR

**Supplemental Figure S5.** (A) Signal intensity comparing across low to high DNA inputs. (B) Spearman's rho for CpG probes rescued by ELBAR (n_diff: number of CpG probes with p-value of ELBAR < 0.05 and p-value of pOOBAH > 0.05). (C – F) ELBAR performance for samples with globally (C) intermediated, (D) fully methylated, (E) lowly methylated, and (F) testicular seminoma tissues.

# S6

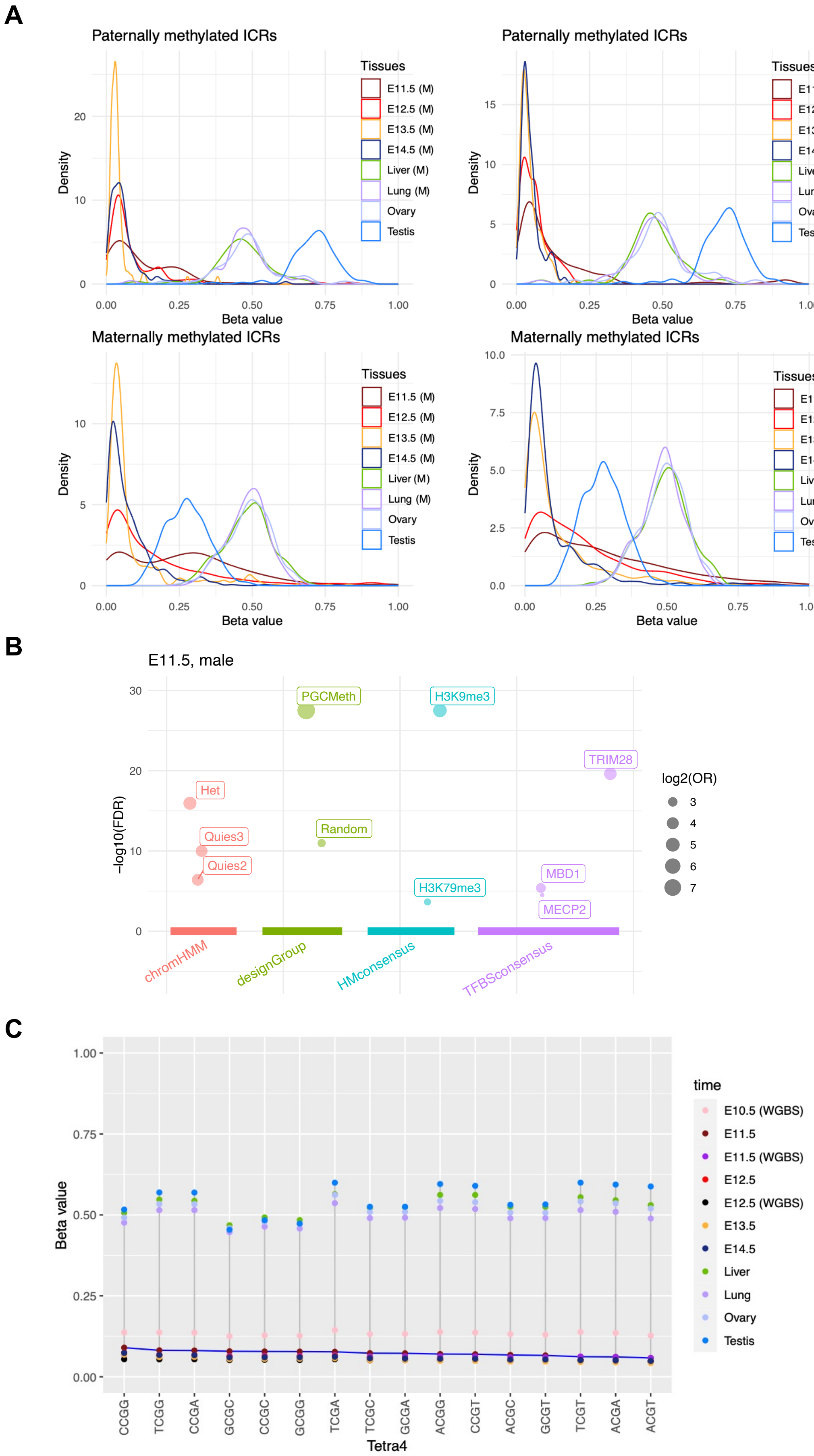
A
Paternally methylated ICRs
Paternally methylated ICRs
Maternally methylated ICRs
Maternally methylated ICRs
Density
Beta value
Tissues
E11.5 (M)
E12.5 (M)
E13.5 (M)
E14.5 (M)
Liver (M)
Lung (M)
Ovary
Testis
E11.5 (F)
E12.5 (F)
E13.5 (F)
E14.5 (F)
Liver (F)
Lung (F)
B
E11.5, male
−log10(FDR)
PGCMeth
H3K9me3
TRIM28
Het
Quies3
Quies2
Random
H3K79me3
MBD1
MECP2
chromHMM
designGroup
HMconsensus
TFBSconsensus
log2(OR)
C
Beta value
Tetra4
CCGG
TCGG
CCGA
GCGC
CCGC
GCGG
TCGA
TCGC
GCGA
ACGG
CCGT
ACGC
GCGT
TCGT
ACGA
ACGT
time
E10.5 (WGBS)
E11.5
E11.5 (WGBS)
E12.5
E12.5 (WGBS)
E13.5
E14.5
Liver
Lung
Ovary
Testis

**Supplemental Figure S6.** (A) DNA methylation distribution in imprinting DMR regions in PGCs and tissues. Figures on the left were generated with CpGs paternally methylated ICRs (CpGs in *Dlk1-Gtl2*, *H19*, and *Rasgrf*), and figures on the right were generated with CpGs maternally methylated ICRs (CpGs in *Gnas1a*, *Grb10*, *Igf2r*, *Impac*t, *Kcnq1ot1*, *Nespas-Gnasxl*, *Peg3*, *Peg10*, *Plagl1*, *Slc38a4*, *Snrpn*, and *Zrsr1*). (B) Highly methylated regions in the four male PGCs of E11.5. Query CpGs are probes with beta values over 0.5 in the samples. (C) Sequence context of loss of DNA methylation in PGCs and tissues. CpGs flanked by A/T or G/C. X-axis is arranged in descending order according to beta values from the E11.5 PGC array data.

### SUPPLEMENTARY TABLES

**Supplemental Table S1.** (A) Experimental workflows analyzed in this study. (B) Methods with different elution size adjustments. (C) Sample data generated by Workflow M. (D) Comparison of preamplification performance from FACS sorted 10 cells to 100 cells of NIH3T3.

**Supplemental Table S2.** (A) Public HM450/EPIC array data used in this study (n=105,475). Information for all Infinium BeadChip data of (B) PGCs produced and (C) tissue samples implemented to compare methylome with PGCs.

### SUPPLEMENTARY METHODS

#### Whole-genome pre-amplification

Workflow F was performed as described in (3) with minor modifications. The 10 μl of bisulfite-converted DNA (BCD) was denatured at 100°C for 5 min and cooled to RT. It was then incubated at 37°C for 2 hours with the five units of Klenow fragment (3'-5' exo-) (NEB, M0212S), 2 μl of 10 X Klenow buffer, 0.4 μl of 10 mM dNTP (BioRad, F560), 4 μl of 10 μM T7-dN6 (5′-GTAATACGACTCACTATAGGGCNNNNNN-3′) primer (IDT) and 2.6 μl of autoclaved ultrapure water. The product was heated at 100 °C again for 5 min and incubated at 37°C for 2 hours with fresh 5 U of Klenow fragment. The reaction was ended by incubation at 75°C for 10 min. Workflow G is performed with the same condition as workflow F but uses 10 units of Klenow fragment. For workflow H, a whole genome amplified product by workflow F was amplified again using workflow G methods. Thus, it achieved a total of 4 cycles of whole genome amplification.

The workflows I and J were performed as described in (4) with minor modifications. 10 μl of BCD was mixed with 0.8 μl of 10 mM dNTP, 4μl of 10 uM of T7-dN6 (workflow I), or random hexamer (NEB, S1230S) (workflow J). The mixture was denatured at 98°C for 1 min and rapidly cooled to ice for 2 min. 10 U of Klenow fragment and 2 μl of 10X Klenow buffer were added and incubated at 4°C 10 min, 4°C/sec to 37°C, 37°C for 30 min. The product was denatured at 98°C for 1 min and rapidly cooled to ice for 2 min. Then, the product was incubated at 4°C for 10 min, 4°C/sec to 37°C, 37°C for 30 min with 0.5 μl of 10X Klenow Buffer, 10 U of Klenow fragment, 0.8 μl of 10 mM dNTP and 1.7 μl of autoclaved ultrapure water. The reaction was ended by incubation at 75°C for 10 min.

Whole genome amplified products were purified using Cytiva Sera-Mag SpeedBeads™ (Fisher, 09-981-123). The bead reagents were prepared as previously described (5). 2x volume of bead reagents were added to the 20 μl of the product, mixed thoroughly, briefly spun down, incubated the mixture for 5 min at room temperature, placed on a magnet stand, and washed 2 times with 500 μl of freshly made 80% EtOH, allowed to dry for 3 to 5 mins. The final elution was performed using 10 – 11 μl autoclaved ultrapure water. 1 μl of the product was used to calculate DNA concentration using Qubit 4.0, and the rest of 10 μl was submitted to microarray analysis.

In workflows K and J, whole-genome amplification was carried out using products of workflows G and I, respectively, with a final elution volume of 20 µl by bead purification. The PCR reaction was performed with purified DNAs, 2.5 µl of 10 µM T7 (5′-GTAATACGACTCACTATAGGGC-3′) primer (IDT), 25 µl of Bioline My Taq Red Mix (Thomas, BIO-25044), and 2.5 µl of autoclaved ultrapure water. The PCR conditions are 72 °C for 10 min, 13 – 26 cycles of 95 °C for 1 min, 60 °C for 1 min, and 72 °C for 2 min. Bead purification of the final product was performed with 11 µl of ultrapure water for elution. 1 µl of the product was used to calculate DNA concentration using Qubit 4.0, and the rest of 10 µl was submitted to microarray analysis.

### SUPPLEMENTARY MATERIAL REFERENCES
